# Comparison of Janus kinase inhibitors to block the type I interferon pathway in human skeletal muscle cells

**DOI:** 10.1101/2021.02.08.430317

**Authors:** Travis B. Kinder, James Inglese

## Abstract

The family of Janus kinases (JAK1, JAK2, JAK3, TYK2) mediate signal transduction from cytokine receptors by phosphorylation and activation of intracellular signaling pathways and transcription factors. Small molecule antagonists of JAKs (Jakinibs) have been developed with varying selectivity for the use in malignancies and immune regulation. There is growing recognition of the effectiveness of jakinibs in autoimmunity of the skeletal muscle called myositis, but which of these drugs is most effective is unknown. We have assayed a library of 48 jakinibs for their ability to inhibit the JAK1/TYK2-dependent type I interferon (IFN) - major histocompatibility complex (MHC) class I pathway using human skeletal muscle cells genome-engineered to fuse a pro-luminescent HiBiT peptide to endogenous MHC class I. The most effective compounds were upadacitinib (JAK1/2 inhibitor, FDA approved) and deucravacitinib (TYK2 inhibitor, phase III). These active jakinibs warrant further clinical evaluation to show their safety and efficacy in patients.

**Graphical Abstract:** 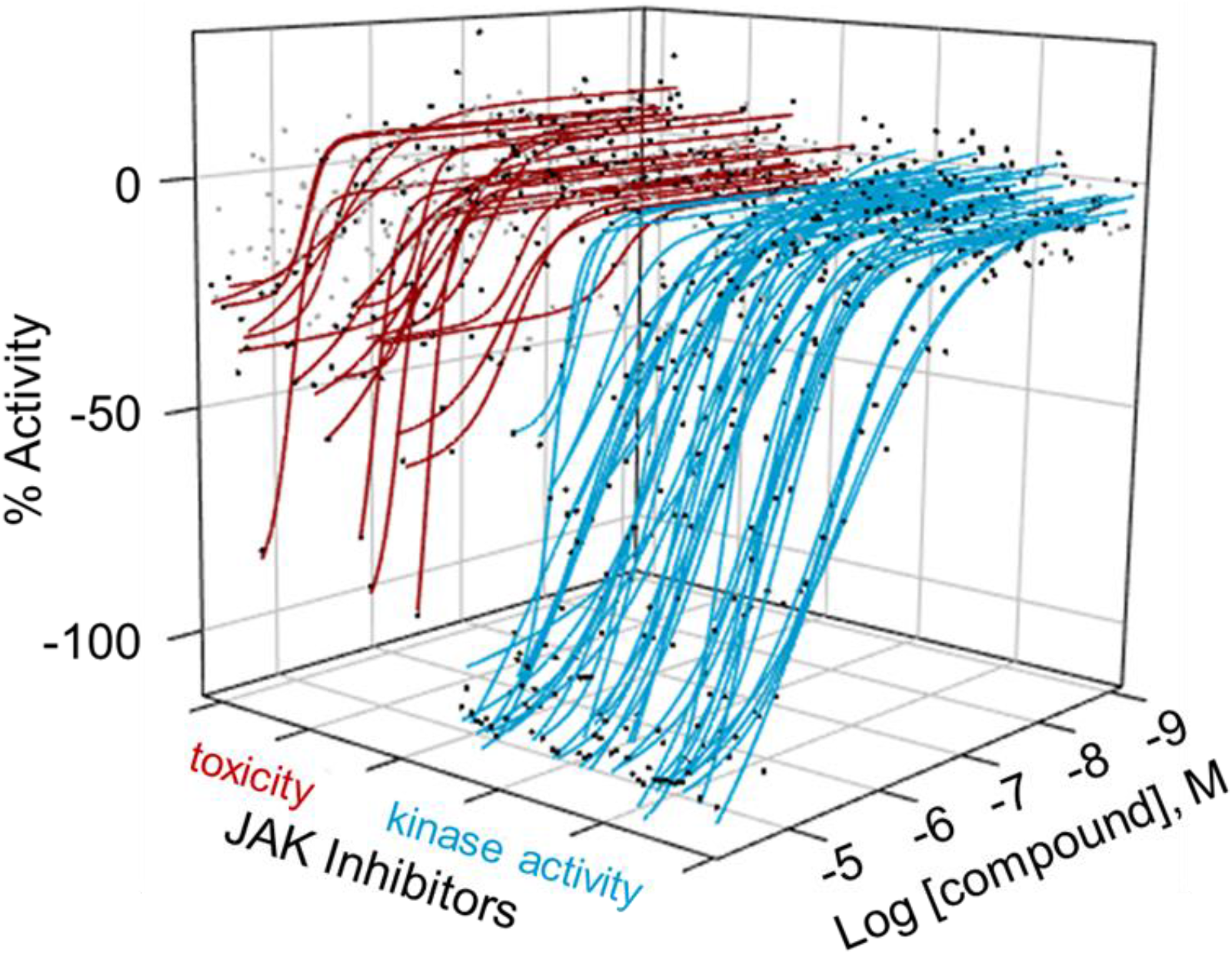

---

Janus kinases comprise four tyrosine kinases JAK1, JAK2, JAK3, and TYK2 that associate with type I/II cytokine receptors to mediate transduction of extracellular signals into alterations in gene expression and cell behavior. Upon ligand binding, cytokine receptors dimerize, and associated JAKs phosphorylate each other and the receptor itself. Activated cytokine receptor/JAK complexes then attract and phosphorylate intracellular signaling mediators such as the signal transducer and activator of transcription proteins (STATs), mitogen-activated protein kinases (MAPK), and phosphoinositide 3-kinase (PI3K).^1^ Phosphorylated STAT transcription factors dimerize, translocate to the nucleus, and promote expression of immune-regulatory genes. STAT-regulated genes are particularly important in autoimmune and autoinflammatory conditions such as the idiopathic inflammatory myopathies (myositis). A subset of myositis called dermatomyositis displays a characteristic fingerprint of up-regulated JAK-STAT mediated genes called the type I IFN signature, which are induced by type I IFN (IFN-α, -β, -ε, -κ, and -ω) binding to IFN-α receptor (IFNAR) and JAK1/TYK2 phosphorylation of STAT1/2.^2^ One such gene induced by type I IFN and thought to play a role in myositis pathology is MHC class I, which presents peptides to T cells, is located within the genomic region most associated with myositis, and its over-expression in mouse skeletal muscle leads to a myositis-like phenotype.^3,4^

Small molecule jakinibs have recently been developed for conditions involving over-active JAK signaling, such as hematologic malignancies, organ transplantation, autoimmunity, or even coronavirus disease 2019 (COVID-19)-associated cytokine storm.^5^ The first of these was a pan-JAK inhibitor tofacitinib approved by the Food and Drug Administration (FDA) in 2012 for rheumatoid arthritis.^6^ Newer generations of compounds have sought various selectivities among the different JAK isoforms to reduce immunosuppression and other side-effects such as decreased hematopoiesis. Small pilot studies and anecdotal evidence exists for the efficacy of jakinibs in myositis, but more studies are needed to determine which are safest and most effective.^7–9^ Described here, we assayed 48 jakinibs in 11-point, 3-fold titrations to measure their cellular potency and efficacy at inhibiting the type I interferon pathway in immortalized human skeletal muscle cell using the HLAB HiBiT cell line we described recently.^10^ Briefly, CRISPR/Cas9 was used to fuse an 11 amino acid pro-luminescent fragment of nanoluciferase (HiBiT) to the C-terminus of endogenous MHC class I (allele HLAB*08:01) in C25cl48 myoblasts. Treatment with recombinant IFN-β increases the expression of HLAB HiBiT fusion as measured by luminescence production after cell lysis and addition of complementary LgBiT peptide and furimazine substrate. Jakinib activity could be assayed by this system through inhibition of the IFNAR-associated JAK1/TYK2 and resulting loss of HiBiT signal compared to IFN-β treatment alone.

## Results and Discussion

A library of 48 jakinibs was compiled through a database search using Palantir software mining PubChem, ChEMBL, DrugBank, and in-house compounds for the terms JAK, JAK1, JAK2, JAK3, and TYK2 within the mechanism of action or target fields. Citeline Informa database showed that as of December 2020, eight of the compounds were approved for use in humans (by either FDA, European Medicines Agency (EMA), or the Japanese Pharmaceuticals and Medical Devices Agency (PMDA)), 13 were in clinical trials, and 27 were preclinical or discontinued. The evidence for targets and selectivity were corroborated through primary literature reviews and included in **Tables 1&2** and **Supplementary Tables 1&2**. A complete reference list for this analysis as well as chemical structures are in **Supplementary Table 2**. We reported the primary target as the kinase most potently inhibited by the compound and off-target kinases as those that were inhibited within 10-fold potency of the primary target. This resulted in a wide variety of kinase inhibitors that were either broadly active, pan-JAK inhibitors, dual JAK inhibitors, and single JAK isoform specific.

**Table 1.** Pharmacologic parameters for approved jakinibs

| Compound | HiBit |  |  |  | CellTiter-Glo / CellTiter-Fluor |  |  |  | Targets | Approval |
| --- | --- | --- | --- | --- | --- | --- | --- | --- | --- | --- |
|  | Curve Class | IC<sub>50</sub> (nM) | Efficacy (%) | Hill Slope | Curve Class | IC<sub>50</sub> (nM) | Cell Death (%) | Hill Slope |  |  |
| Upadacitinib | -1a | 330 ± 62 | -98 ± 6 | 0.83 ± 0.23 | -1b | 8,800 ± 2,500 | -30 ± 3 | 4.17 ± 0.99 | **JAK1/2** | FDA |
| Ruxolitinib | -1a | 860 ± 58 | -99 ± 5 | 1.03 ± 0.12 | 4 |  |  |  | **JAK1/2/TYK2** | FDA |
| Baricitinib | -1a | 1,300 ± 250 | -96 ± 5 | 0.87 ± 0.12 | 4 |  |  |  | **JAK1/2/TYK2** | FDA |
| Peficitinib | -1a | 1,700 ± 120 | -95 ± 7 | 1.00 ± 0.10 | 4 |  |  |  | JAK1/2/**3**/TYK2, others? | PMDA |
| Tofacitinib | -1a | 2,200 ± 500 | -95 ± 7 | 0.83 ± 0.06 | 4 |  |  |  | **JAK1/2/3** | FDA |
| Fedratinib | -1a | 3,600 ± 490 | -99 ± 5 | 1.57 ± 0.41 | -3 | 18,000 ± 1,200 | -70 ± 7 |  | **JAK2**, FLT3 | FDA |
| Filgotinib | -2a | 15,300 ± 1,000 | -100 ± 4 | 1.17 ± 0.24 | -3 |  | -31 ± 4 |  | **JAK1/2/TYK2** | EMA, PMDA |
| Leflunomide | -2b | 17,000 ± 1,200 | -70 ± 5 | 1.21 ± 0.41 | -3 |  | -14 ± 8 |  | **DHODH**, JAK1/3, LCK, FYN | FDA |

IC<sub>50</sub> - half maximal inhibitory concentration, primary targets are in bold red, CellTiter-Glo data shown for all but CellTiter-Fluor data shown for Fedratinib

**Table 2.**
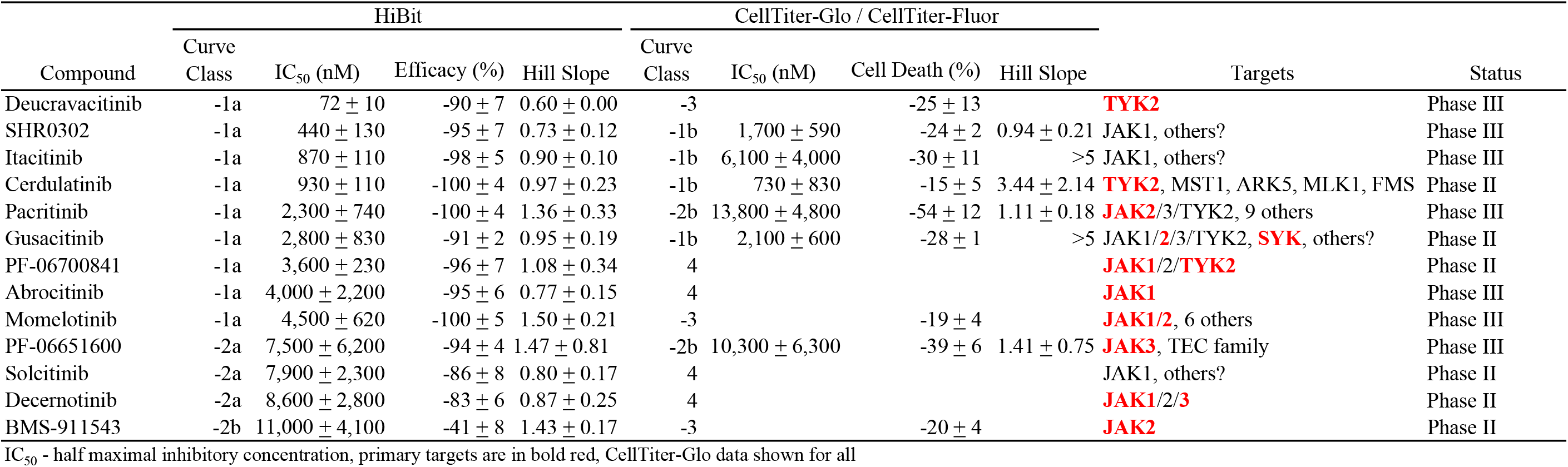
Pharmacologic parameters for jakinibs in clinical trials

Compound titrations were assayed for efficacy and potency using HLAB HiBiT cells in the presence of IFN-β, and cell viability was determined using the same cells and conditions but assayed by CellTiter-Glo (CTG, measure of cellular ATP) and CellTiter-Fluor (CTF, measures live-cell protease activity). CTG resulted in the most robust and clean viability data, but yielded inverse bell-shaped curves for 4 compounds (fedratinib, lestauritinib, RO495, and curcubitacin I), for which CTF yielded standard loss-of-signal curves. For compounds with bell-shaped CTG curves, both CTG and CTF assay results were presented. All assays were run in triplicate independent experiments on different days. The resulting concentration-response curves (CRCs) for approved drugs are shown in **Figure 1**, clinical candidates in **Figure 2**, and preclinical/discontinued compounds in **Supplementary Figure 1**. CRCs were categorized into classes where curve class 1 was the most potent (complete curve with both asymptotes), class 2 was less potent (an incomplete curve with one asymptote), class 3 had a single point activity, and class 4 was inactive. Additionally, an “a” or “b” suffix represents a high efficacy of ≥80% or a lower efficacy of <80%, respectively, and negative curves were inhibitory while positive were stimulatory.^11^ The curve classes, measurements for potency, efficacy, and viability for approved drugs are shown in **Table 1**, clinical candidates in **Table 2**, and preclinical/discontinued compounds in **Supplementary Table 1**. As these compounds had known jakinib activity, unsurprisingly, most were fairly potent and efficacious in this assay, with twenty-eight compounds in curve class -1a, ten in curve class -2a, nine in curve class -2b, and three were inactive class 4.

**Figure 1.**
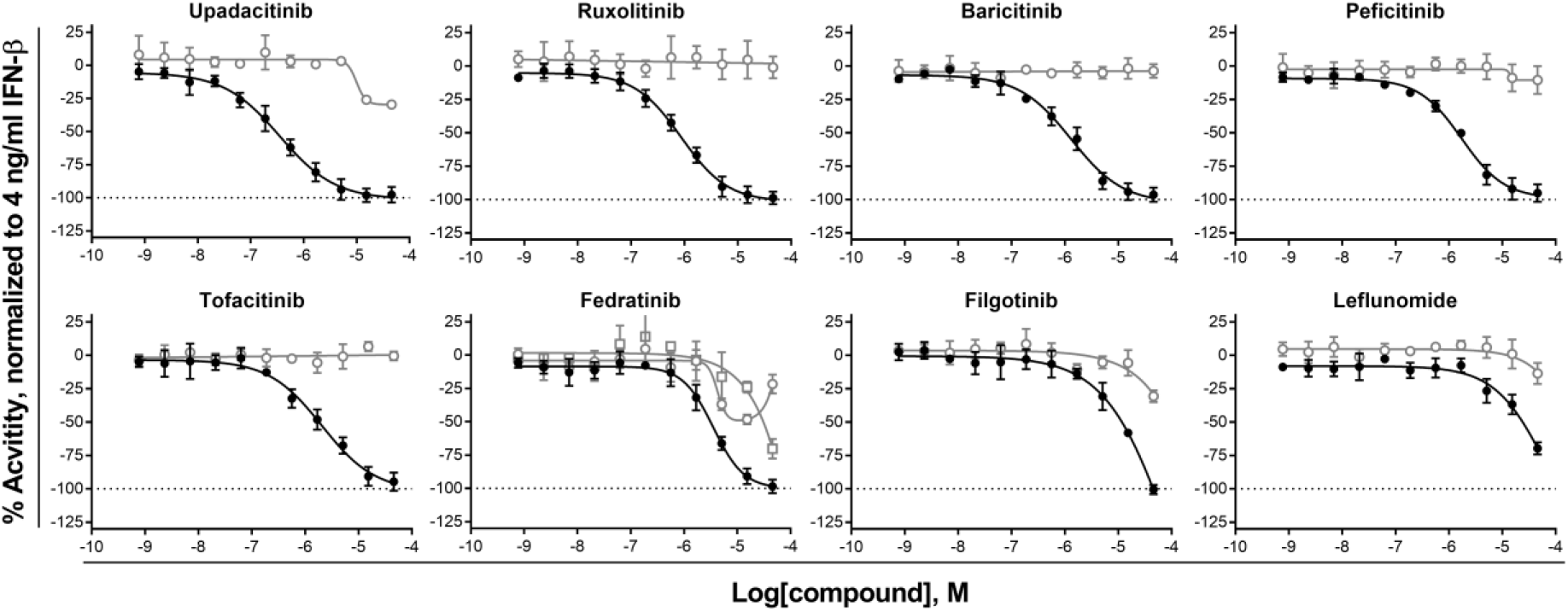
CRCs for approved jakinibs. HLA-B HiBit clone AB-5 cells plated in 1536 well plates and treated for 24 hours with 4.0 ng/ml IFN-β plus 11-point, 1:3 titrations of compounds. Black circles – HiBit, open gray circles – CTG, open gray squares - CTF. For HiBit efficacy assays, data was normalized within each plate to 4.0 ng/ml IFN-β = 0% and DMSO alone = -100% activity. For CTG and CTF cytotoxicity assays, data was normalized within each plate to 4.0 ng/ml IFN-β = 0% and 92 uM Digitonin = -100% activity.

**Figure 2.**
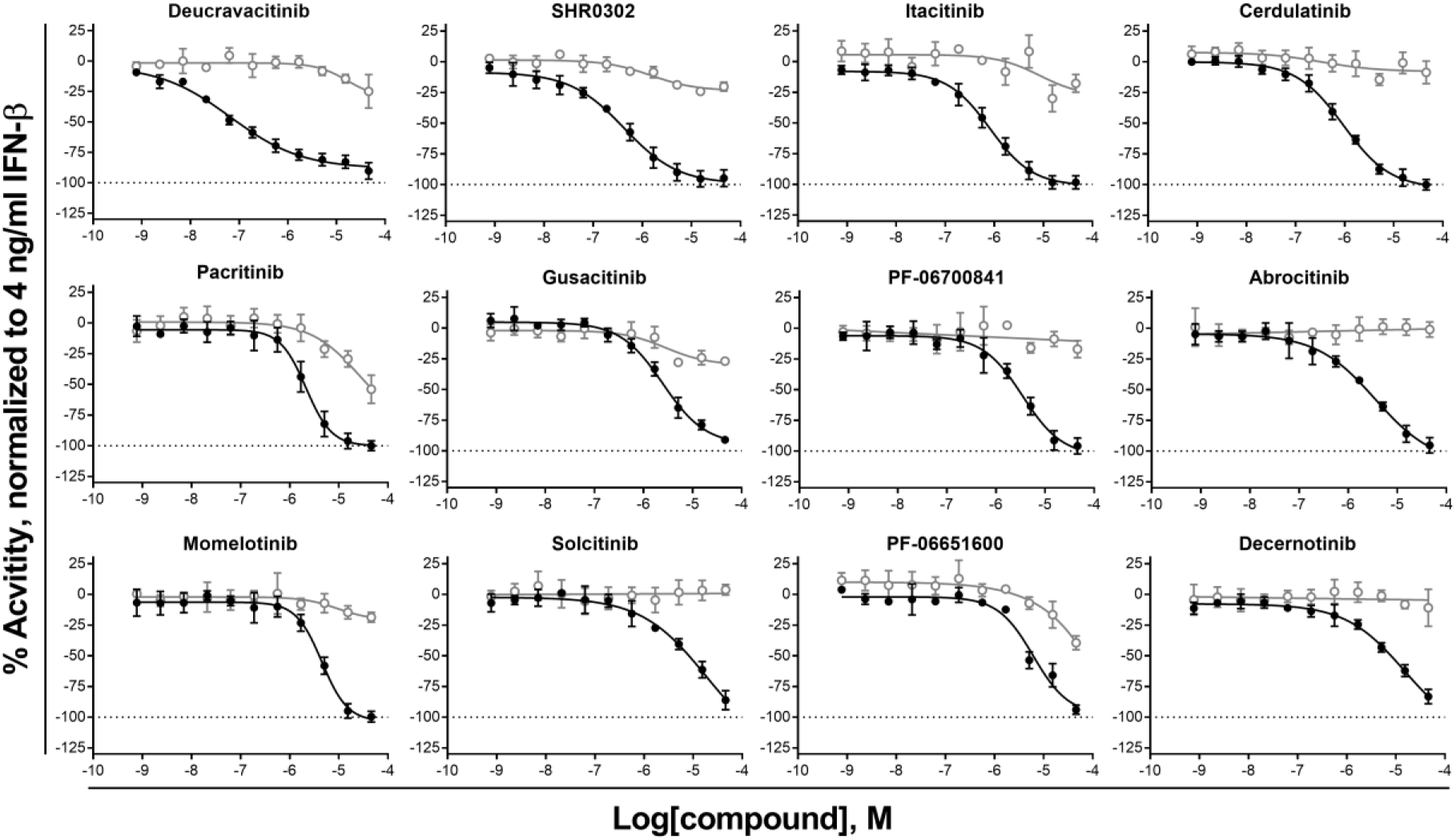
CRCs for jakinibs in clinical trials. HLA-B HiBit clone AB-5 cells plated in 1536 well plates and treated for 24 hours with 4.0 ng/ml IFN-β plus 11-point, 1:3 titrations of compounds. Black circles – HiBit, open gray circles – CTG, open gray squares - CTF. For HiBit efficacy assays, data was normalized within each plate to 4.0 ng/ml IFN-β = 0% and DMSO alone = -100% activity. For CTG and CTF cytotoxicity assays, data was normalized within each plate to 4.0 ng/ml IFN-β = 0% and 92 uM Digitonin = -100% activity.

The approved drugs upadacitinib, ruxolitinib, baricitinib, peficitinib, and tofacitinib had 100% efficacy, little to no cell death, and ranged in potencies between the most active upadacitinib IC_50_ = 330 ± 62 nM to the least active tofacitinib IC_50_ = 2,200 ± 500 nM. Fedratinib is a JAK2/FLT3 inhibitor that appeared to have a -1a curve class, but was confounded by a significant reduction in cell viability under the effective concentrations, which may be due to its FLT3 inhibition.^12^ Filgotinib, a JAK1/2/TYK2 inhibitor approved by EMA and PMDA, was not very active here (curve class -2a) and had some toxicity at the higher concentrations. Finally, Leflunomide was least active of the approved drugs (curve class -2b) with some toxicity, and is actually an antirheumatic drug with the primary target being dihydroorotate dehydrogenase (DHODH), but also inhibits JAKs and other kinases at higher concentrations.^13^

The clinical candidate deucravacitinib was the most efficacious and potent compound in this library with 100% efficacy and IC_50_ = 72 ± 10 nM. Deucravacitinib is a highly specific TYK2 inhibitor in phase III clinical trials for psoriasis as well as phase II trials for psoriatic arthritis, inflammatory bowel disease, and lupus. This compound is novel in that it binds to an allosteric site on the JH2 pseudokinase domain of TYK2, whereas kinase inhibitors generally competitively target the ATP binding site.^14^ Targeting the ATP binding site with traditional kinase inhibitors, including the approved jakinibs, is challenging because 1) the compound needs to outcompete ATP which is at high concentrations inside the cell, and 2) the ATP binding site is highly conserved across the kinome, making it difficult to achieve selectivity toward just one kinase.

The next five most potent clinical candidates (SHR0302, itacitinib, cerdulatinib, pacritinib, and gusacitinib) had toxicity curve IC_50_s within 10-fold of the HiBiT curve IC_50_s, meaning that the compounds were killing cells to some extent within the effective concentration range. However, at the highest tested concentration (50 μM), they ranged from mildly toxic with cerdulatinib cell death at 15% to moderately toxic with pacritnib killing 54% of cells. The remaining seven clinical candidates were not potent, having IC_50_ values > 3,600 nM.

The preclinical/discontinued compounds were generally weak or had toxicity. Many were older molecular probes that never got approval or were discontinued due to adverse events. The side-effects and lack of translation in the clinic may be due to off-target kinase activity, as shown in cell-free kinase panel screening for many of these compounds. For example, lestauritinib is a synthetic analogue of the pan-kinase inhibitor staurosporine that binds to 115 kinases within 10-fold of the K_D_ for JAKs, and is most potent towards phosphorylase b kinase gamma catalytic chain.^15^ Therefore lestauritinib is a JAK inhibitor, but JAKs are not the primary target and it inhibits many other kinases.

It has recently become clear that clinical advancement of selective kinase inhibitors should include screening against a large panel of purified kinases representing a broad array of the human kinome (estimated 518 kinases) as well as phenotypic cell-based models to determine selectivity.^16^ It is well known that kinase inhibitors can promiscuously inhibit off-target enzymes, which may be a source of side-effects in clinical use.^15^ For compounds that did not have published data from kinome panel screening, the *targets* column in **Tables 1&2** and **Supplementary Tables 1&2** listed the JAKs inhibited followed by the phrase “others?” In addition, assays should be carried out at physiological concentrations of ATP (1-5 mM) in order to accurately reflect the selectivity of compounds in a biologically relevant context.^17^ Not all compounds here have been screened through such panels, and if they have, their methods varied greatly. This makes it difficult to calculate selectivity scores that are comparable between studies. In general though, the approved drugs in this study inhibited the fewest number of off-target kinases compared to clinical trial, preclinical, and discontinued compounds. In addition, compounds that had more off-target kinases showed more toxicity.

Overall, the two most potent compounds in this library for inhibiting the type I IFN pathway in skeletal muscle cells with least cell death was upadacitinib (JAK1/2 inhibitor, FDA approved) and deucravacitinib (TYK2 inhibitor, phase III). This makes sense as the two kinases that mediate the type I IFN pathway are JAK1 and TYK2.^1^ Although upadacitinib is listed here as a JAK1/2 inhibitor, it is reported to have 3-fold selectivity for JAK1 over JAK2, but would still result in significant JAK2 inhibition at concentrations that fully inhibit JAK1. These results warrant further investigations into the usefulness of these drugs in myositis animal models or patients, especially since there exists clinical data in autoimmunity for both. Reported here is a simplified model that only includes muscle cells and the type I IFN pathway, so more data should be taken into consideration when comparing different jakinibs and assessing their efficacy and safety in patients. For example, immunosuppression with a more broadly acting jakinibs maybe a benefit or side-effect to patients depending on the level of involvement of different JAK-STAT pathways and different cell types.

## Methods

### Cell culture, HiBiT, and CTG assays

The methods used for cell culture, compound treatment, HiBiT assay, and CTG assay were the same as described previously,^10^ with the exception that here HLAB HiBiT clone AB5 was used instead of AB7 and at a slightly lower cell number of 250 cells/well. Clone AB7 had small genomic duplications surrounding the HiBiT sequence, whereas AB5 had correct HiBiT integration without such aberrations.

### CellTiter-Flour

Gly-Phe-7-Amino-4-Trifluoromethylcoumarin (GF-AFC, MP biomedicals, SKU 03AFC03325) substrate was diluted in PBS and 1.0 μL/well was added with a Multidrop Combi Reagent Dispenser (Thermo Fisher) to a final concentration of 25 uM. Plates were incubated at 37 °C, 5% CO_2_, 95% relative humidity for 1:15 hr, and fluorescence was measured with an Envision 2104 Multilabel Plate Reader (PerkinElmer), Wallac EnVision Manager software v1.12, top read using LANCE/DELFIA mirror, MOCAc 400 excitation filter, and FITC 535 emission filter.

## Acknowledgements

We are appreciative of K. Wilson, PhD (NCATS) for training and assistance with Palantir database searches, the compound management team (NCATS) for preparing the library plate, and P. K. Dranchak, PhD and B. Queme (NCATS) for generation of the 3-axes plot graphical abstract.

## References

(1) González-Navajas, J. M.; Lee, J.; David, M.; Raz, E. Immunomodulatory Functions of Type I Interferons. Nat. Rev. Immunol. 2012, 12 (2), 125–135. https://doi.org/10.1038/nri3133.

(2) Gallay, L.; Mouchiroud, G.; Chazaud, B. Interferon-Signature in Idiopathic Inflammatory Myopathies. Curr. Opin. Rheumatol. 2019, 31 (6), 634–642. https://doi.org/10.1097/BOR.0000000000000653.

(3) Miller, F. W.; Lamb, J. A.; Schmidt, J.; Nagaraju, K. Risk Factors and Disease Mechanisms in Myositis. Nat. Rev. Rheumatol. 2018, 14 (5), 255–268. https://doi.org/10.1038/nrrheum.2018.48.

(4) Nagaraju, K.; Casciola-Rosen, L.; Lundberg, I.; Rawat, R.; Cutting, S.; Thapliyal, R.; Chang, J.; Dwivedi, S.; Mitsak, M.; Chen, Y.-W.; Plotz, P.; Rosen, A.; Hoffman, E.; Raben, N. Activation of the Endoplasmic Reticulum Stress Response in Autoimmune Myositis: Potential Role in Muscle Fiber Damage and Dysfunction. Arthritis Rheum. 2005, 52 (6), 1824–1835. https://doi.org/10.1002/art.21103.

(5) Valenzuela-Almada, M. O.; Putman, M. S.; Duarte-García, A. The Protective Effect of Rheumatic Disease Agents in COVID-19. Best Pract. Res. Clin. Rheumatol. 2021, 101659. https://doi.org/10.1016/j.berh.2021.101659.

(6) Changelian, P. S.; Flanagan, M. E.; Ball, D. J.; Kent, C. R.; Magnuson, K. S.; Martin, W. H.; Rizzuti, B. J.; Sawyer, P. S.; Perry, B. D.; Brissette, W. H.; McCurdy, S. P.; Kudlacz, E. M.; Conklyn, M. J.; Elliott, E. A.; Koslov, E. R.; Fisher, M. B.; Strelevitz, T. J.; Yoon, K.; Whipple, D. A.; Sun, J.; Munchhof, M. J.; Doty, J. L.; Casavant, J. M.; Blumenkopf, T. A.; Hines, M.; Brown, M. F.; Lillie, B. M.; Subramanyam, C.; Shang-Poa, C.; Milici, A. J.; Beckius, G. E.; Moyer, J. D.; Su, C.; Woodworth, T. G.; Gaweco, A. S.; Beals, C. R.; Littman, B. H.; Fisher, D. A.; Smith, J. F.; Zagouras, P.; Magna, H. A.; Saltarelli, M. J.; Johnson, K. S.; Nelms, L. F.; Des Etages, S. G.; Hayes, L. S.; Kawabata, T. T.; Finco-Kent, D.; Baker, D. L.; Larson, M.; Si, M. S.; Paniagua, R.; Higgins, J.; Holm, B.; Reitz, B.; Zhou, Y. J.; Morris, R. E.; O’Shea, J. J.; Borie, D. C. Prevention of Organ Allograft Rejection by a Specific Janus Kinase 3 Inhibitor. Science (80-.). 2003, 302 (5646), 875–878. https://doi.org/10.1126/science.1087061.

(7) Kim, H.; Dill, S.; O’Brien, M.; Vian, L.; Li, X.; Manukyan, M.; Jain, M.; Adeojo, L. W.; George, J.; Perez, M.; Grom, A. A.; Sutter, M.; Feldman, B. M.; Yao, L.; Millwood, M.; Brundidge, A.; Pichard, D. C.; Cowen, E. W.; Shi, Y.; Lu, S.; Tsai, W. L.; Gadina, M.; Rider, L. G.; Colbert, R. A. Janus Kinase (JAK) Inhibition with Baricitinib in Refractory Juvenile Dermatomyositis. Ann. Rheum. Dis. 2020, annrheumdis-2020-218690. https://doi.org/10.1136/annrheumdis-2020-218690.

(8) Ladislau, L.; Suárez-Calvet, X.; Toquet, S.; Landon-Cardinal, O.; Amelin, D.; Depp, M.; Rodero, M. P.; Hathazi, D.; Duffy, D.; Bondet, V.; Preusse, C.; Bienvenu, B.; Rozenberg, F.; Roos, A.; Benjamim, C. F.; Gallardo, E.; Illa, I.; Mouly, V.; Stenzel, W.; Butler-Browne, G.; Benveniste, O.; Allenbach, Y. JAK Inhibitor Improves Type I Interferon Induced Damage: Proof of Concept in Dermatomyositis. Brain 2018, 141 (6), 1609–1621. https://doi.org/10.1093/brain/awy105.

(9) Paik, J.; Albayda, J.; Tiniakou, E.; Koenig, A. Study of Tofacitinib in Refractory Dermatomyositis (STIR): An Open Label Pilot Study in Refractory Dermatomyositis - ACR Meeting Abstracts. Arthritis Rheum 2018, 70, L02.

(10) Kinder, T. B.; Dranchak, P. K.; Inglese, J. High-Throughput Screening to Identify Inhibitors of the Type I Interferon–Major Histocompatibility Complex Class I Pathway in Skeletal Muscle. ACS Chem. Biol. 2020, 15 (7), 1974–1986. https://doi.org/10.1021/acschembio.0c00343.

(11) Inglese, J.; Auld, D. S.; Jadhav, A.; Johnson, R. L.; Simeonov, A.; Yasgar, A.; Zheng, W.; Austin, C. P. Quantitative High-Throughput Screening: A Titration-Based Approach That Efficiently Identifies Biological Activities in Large Chemical Libraries. Proc. Natl. Acad. Sci. U. S. A. 2006, 103 (7), 11473–11478. https://doi.org/10.1073/pnas.0604348103.

(12) Wernig, G.; Kharas, M. G.; Okabe, R.; Moore, S. A.; Leeman, D. S.; Cullen, D. E.; Gozo, M.; McDowell, E. P.; Levine, R. L.; Doukas, J.; Mak, C. C.; Noronha, G.; Martin, M.; Ko, Y. D.; Lee, B. H.; Soll, R. M.; Tefferi, A.; Hood, J. D.; Gilliland, D. G. Efficacy of TG101348, a Selective JAK2 Inhibitor, in Treatment of a Murine Model of JAK2V617F-Induced Polycythemia Vera. Cancer Cell 2008, 13 (4), 311–320. https://doi.org/10.1016/j.ccr.2008.02.009.

(13) Elder, R. T.; Xu, X.; Williams, J. W.; Gong, H.; Finnegan, A.; Chong, A. S. F. The Immunosuppressive Metabolite of Leflunomide, A77 1726, Affects Murine T Cells Through Two Biochemical Mechanisms. J. Immunol. 1997, 159 (1), 22–27.

(14) Burke, J. R.; Cheng, L.; Gillooly, K. M.; Strnad, J.; Zupa-Fernandez, A.; Catlett, I. M.; Zhang, Y.; Heimrich, E. M.; McIntyre, K. W.; Cunningham, M. D.; Carman, J. A.; Zhou, X.; Banas, D.; Chaudhry, C.; Li, S.; D’Arienzo, C.; Chimalakonda, A.; Yang, X. X.; Xie, J. H.; Pang, J.; Zhao, Q.; Rose, S. M.; Huang, J.; Moslin, R. M.; Wrobleski, S. T.; Weinstein, D. S.; Salter-Cid, L. M. Autoimmune Pathways in Mice and Humans Are Blocked by Pharmacological Stabilization of the TYK2 Pseudokinase Domain. Sci. Transl. Med. 2019, 11 (502). https://doi.org/10.1126/scitranslmed.aaw1736.

(15) Davis, M. I.; Hunt, J. P.; Herrgard, S.; Ciceri, P.; Wodicka, L. M.; Pallares, G.; Hocker, M.; Treiber, D. K.; Zarrinkar, P. P. Comprehensive Analysis of Kinase Inhibitor Selectivity. Nat. Biotechnol. 2011, 29 (11), 1046–1051. https://doi.org/10.1038/nbt.1990.

(16) Uitdehaag, J. C. M.; Verkaar, F.; Alwan, H.; De Man, J.; Buijsman, R. C.; Zaman, G. J. R. A Guide to Picking the Most Selective Kinase Inhibitor Tool Compounds for Pharmacological Validation of Drug Targets. British Journal of Pharmacology. Br J Pharmacol June 2012, pp 858–876. https://doi.org/10.1111/j.1476-5381.2012.01859.x.

(17) Thorarensen, A.; Banker, M. E.; Fensome, A.; Telliez, J. B.; Juba, B.; Vincent, F.; Czerwinski, R. M.; Casimiro-Garcia, A. ATP-Mediated Kinome Selectivity: The Missing Link in Understanding the Contribution of Individual JAK Kinase Isoforms to Cellular Signaling. ACS Chem. Biol. 2014, 9 (7), 1552–1558. https://doi.org/10.1021/cb5002125.

