## Supporting Information for "Comparison of Janus kinase inhibitors to block the type I interferon pathway in human skeletal muscle cells"

**Supplemental Table 1.** Pharmacologic parameters for preclinical and discontinued jakinibs

| | HiBit | | | | CellTiter-Glo / CellTiter-Fluor | | | | | |
|---|---|---|---|---|---|---|---|---|---|---|
| Compound | Curve class | $IC_{50}$ (nM) | Efficacy (%) | Hill Slope | Curve class | $IC_{50}$ (nM) | Cell Death (%) | Hill Slope | Targets | Status |
| CHZ868 | -1a | 385 ± 130 | -98 ± 5 | 0.94 ± 0.16 | -1b | 230 ± 110 | -42 ± 2 | 1.29 ± 0.52 | **PDGFR**, VEGFR, 11 others, JAK2/TYK2 | Preclinical |
| Lestaurtinib | -1a | 420 ± 48 | -100 ± 4 | 0.63 ± 0.06 | -1b | 68 ± 22 | -19 ± 3 | 2.74 ± 2.39 | **PHKG1**, JAK1/2/3/TYK2, 114 others | Discontinued |
| AZ-960 | -1a | 1,100 ± 450 | -99 ± 5 | 1.14 ± 0.12 | -1b | 2,400 ± 1,500 | -39 ± 9 | 2.36 ± 1.55 | JAK1/**2**/3/TYK2, TrkA, AURKA, ARK5 | Preclinical |
| Pyridone 6 | -1a | 1,300 ± 220 | -94 ± 6 | 1.18 ± 0.07 | -1b | 1,900 ± 350 | -18 ± 7 | 2.05 ± 0.88 | **JAK1**/2/3/**TYK2**, 25 others | Preclinical |
| AZD-1480 | -1a | 1,500 ± 310 | -97 ± 6 | 1.12 ± 0.24 | -1b | 4,000 ± 260 | -57 ± 4 | 2.68 ± 2.00 | **TrkA**, JAK1/2, AURKA, FLT4, FGFR1, AX1, ARK5 | Discontinued |
| RO495 | -1a | 2,300 ± 900 | -101 ± 4 | 1.21 ± 0.44 | -3 | 12,000 ± 12,000 | -30 ± 3 | | TYK2, others? | Preclinical |
| TG-101209 | -1a | 2,500 ± 560 | -96 ± 7 | 1.11 ± 0.22 | -1b | 1,600 ± 270 | -35 ± 4 | 2.52 ± 0.47 | **JAK2**, FLT3/4, RET, ABL1, FGR, LCK, BRD4 | Preclinical |
| CEP-33779 | -1a | 3,100 ± 560 | -97 ± 6 | 1.15 ± 0.31 | 4 | | | | **JAK2** | Preclinical |
| GDC-046 | -1a | 4,500 ± 290 | -98 ± 5 | 1.70 ± 0.28 | 4 | | | | TYK2, others? | Preclinical |
| Oclacitinib | -1a | 4,700 ± 880 | -96 ± 6 | 0.87 ± 0.12 | 4 | | | | **JAK1/2**/3/TYK2 | Vetrinary |
| SAR-20347 | -2a | 5,300 ± 990 | -100 ± 5 | 0.90 ± 0.20 | -1b | 5,500 ± 1,200 | -37 ± 6 | 3.33 ± 1.04 | **TYK2** | Preclinical |
| TG-46 | -2a | 5,500 ± 1,200 | -94 ± 4 | 1.15 ± 0.18 | -1b | 7,500 ± 1,400 | -49 ± 2 | 1.56 ± 0.61 | JAK2, FLT3, RET, others? | Preclinical |
| Cucurbitacin I | -1a | 5,700 ± 370 | -102 ± 3 | 1.01 ± 0.36 | -2a | 4,300 ± 3,300 | -86 ± 30 | 0.85 ± 0.43 | JAK2, others? | Preclinical |
| NCGC00244250 | -1a | 6,600 ± 760 | -99 ± 5 | 1.36 ± 0.11 | -3 | | -11 ± 2 | | JAK1/TYK2, others? | Preclinical |
| Degrasyn | -1a | 7,400 ± 0 | -102 ± 3 | 2.91 ± 1.39 | -3 | | -81 ± 6 | | **DUBs** - downregulates JAK2, BCR/ABL, MYC | Discontinued |
| TG-89 | -2a | 10,800 ± 5,100 | -95 ± 5 | 1.03 ± 0.06 | -1b | 7,200 ± 1,200 | -41 ± 2 | >5 | JAK2, FLT3, RET, others? | Preclinical |
| Bayer-18 | -2b | 12,000 ± 5,400 | -44 ± 4 | 1.29 ± 0.37 | 4 | | | | TYK2, others? | Preclinical |
| ZM-39923 | -2a | 14,000 ± 930 | -101 ± 3 | 2.02 ± 0.35 | -3 | | -74 ± 18 | | **TGM2**, JAK1/3, EGFR | Preclinical |
| Gandotinib | -2a | 14,000 ± 930 | -96 ± 5 | 0.80 ± 0.00 | -1b | 12,000 ± 1,400 | -23 ± 5 | >5 | JAK1/**2**, FLT3 | Discontinued |
| JAK3i | -2b | 17,900 ± 5,900 | -39 ± 6 | 2.28 ± 1.08 | 4 | | | | **JAK3**, FLT3, TEC-family | Preclinical |
| NVP-BSK805 | -2a | 18,000 ± 1,200 | -99 ± 4 | 1.24 ± 0.26 | -3 | | -76 ± 6 | | **JAK2** | Preclinical |
| XL019 | -2b | 18,000 ± 3,900 | -50 ± 2 | 2.85 ± 1.09 | 4 | | | | **JAK2** | Preclinical |
| JAK2 inhibitor 13 | -2b | 19,000 ± 0 | -50 ± 5 | 1.48 ± 0.67 | 1b | 990 ± 1,700 | -11 ± 9 | 1.21 ± 0.54 | JAK2, others? | Preclinical |
| TCS 21311 | -2b | 20,000 ± 2,800 | -48 ± 6 | 3.56 ± 0.52 | -3 | | -11 ± 5 | | **JAK3**, GSK-3β, PKCα and PKCθ | Preclinical |
| Curvularin | 4 | | | | 4 | | | | JAK1/2, others? | Preclinical |
| JANEX-1 | 4 | | | | 4 | | | | **JAK3** | Preclinical |
| AG 490 | 4 | | | | 4 | | | | **JAK2**/3, EGFR | Discontinued |

$IC_{50}$ - half maximal inhibitory concentration, primary targets are in bold red, CellTiter-Glo data shown for all but CellTiter-Fluor data shown for lestauritinib, RO495, and curcubitacin I

**Supplementay Table 2**. Jakinib structures and references

| Compound | Structure | Status | Targets | References |
|---|---|---|---|---|
| Upadacitinib | 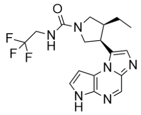 | FDA | **JAK1**/2 | 1 |
| Ruxolitinib | 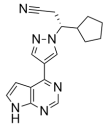 | FDA | **JAK1/2**/TYK2 | 2 |
| Baricitinib | 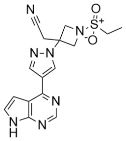 | FDA | **JAK1/2**/TYK2 | 3 |
| Peficitinib | 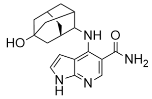 | PMDA | JAK1/2/**3**/TYK2, others? | 4 |
| Tofacitinib | 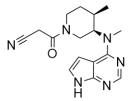 | FDA | **JAK1**/2/3 | 2 |
| Fedratinib | 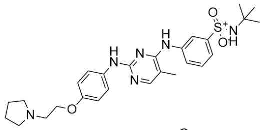 | FDA | **JAK2**, FLT3 | 5 |
| Filgotinib | 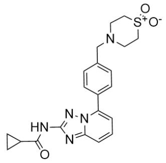 | EMA, PMDA | **JAK1**/2/TYK2 | 2 |
| Leflunomide | 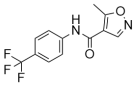 | FDA | **DHODH**, JAK1/3, LCK, FYN | 6 |
| Deucravacitinib | 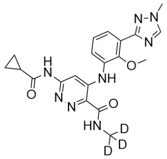 | Phase III | **TYK2** | 7 |
| SHR0302 | 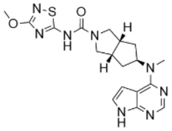 | Phase III | JAK1, others? | 8 |
| Itacitinib | 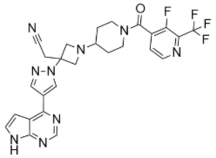 | Phase III | JAK1, others? | 9 |
| Cerdulatinib | 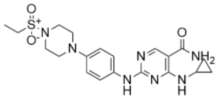 | Phase II | **TYK2**, MST1, ARK5, MLK1, FMS | 10 |
| Pacritinib | 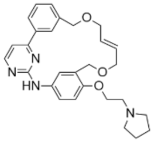 | Phase III | **JAK2**/3/TYK2, 9 others | 11 |
| Gusacitinib | 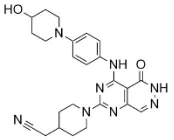 | Phase II | JAK1/2/3/TYK2, SYK, others? | 12 |
| PF-06700841 | 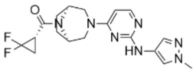 | Phase II | **JAK1**/2/**TYK2** | 13 |
| Abrocitinib | 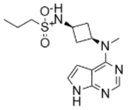 | Phase III | **JAK1** | 14 |
| Momelotinib | 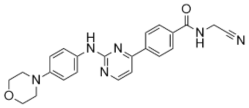 | Phase III | **JAK1/2**, 6 others | 15 |
| PF-06651600 | 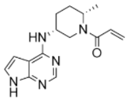 | Phase III | **JAK3**, TEC family | 16 |

| Compound | Structure | Status | Targets | References |
|---|---|---|---|---|
| Solcitinib | 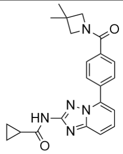 | Phase II | JAK1, others? | 17 |
| Decernotinib | 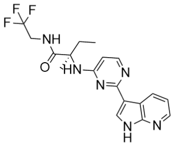 | Phase II | **JAK1**/2/**3** | 2 |
| BMS-911543 | 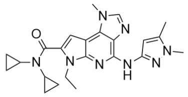 | Phase II | **JAK2** | 18 |
| CHZ868 | 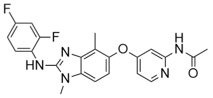 | Preclinical | **PDGFR**, 12 others, JAK2/TYK2 | 19 |
| Lestaurtinib | 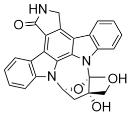 | Discontinued | **PHKG1**, JAKs, 114 others | 20 |
| AZ-960 | 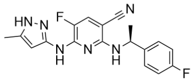 | Preclinical | JAK1/**2**/3/TYK2, TrkA, AURKA, ARK5 | 21 |
| Pyridone 6 | 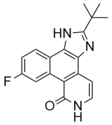 | Preclinical | **JAK1**/2/3/**TYK2**, 25 others | 22, 23 |
| AZD-1480 | 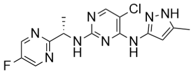 | Discontinued | **TrkA**, JAK1/2, AURKA, FLT4, FGFR1, AX1, ARK5 | 24 |
| RO495 | 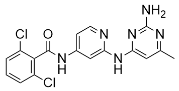 | Preclinical | TYK2, others? | 25 |
| TG-101209 | 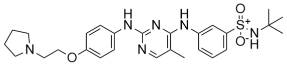 | Preclinical | **JAK2**, FLT3/4, RET, ABL1, FGR, LCK, BRD4 | 26, 27 |
| CEP-33779 | 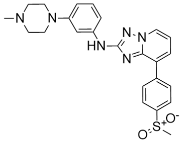 | Preclinical | **JAK2** | 28 |
| GDC-046 | 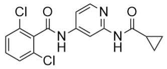 | Preclinical | TYK2, others? | 25 |
| Oclacitinib | 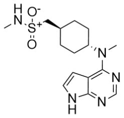 | Veterinary | **JAK1/2**/3/TYK2 | 29 |
| SAR-20347 | 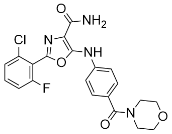 | Preclinical | **TYK2** | 30 |
| TG-46 | 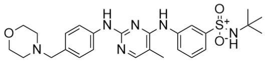 | Preclinical | JAK2, FLT3, RET, others? | 31 |
| Cucurbitacin I | 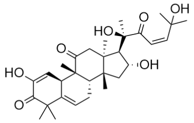 | Preclinical | JAK2, others? | 32 |
| NCGC00244250 | 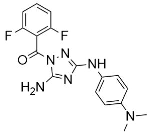 | Preclinical | JAK1/TYK2, others? | 33 |
| Degrasyn | 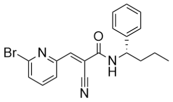 | Discontinued | **Deubiquitinases** | 34 |

| Compound | Structure | Status | Targets | References |
|---|---|---|---|---|
| TG-89 | 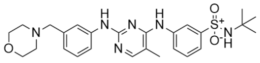 | Preclinical | JAK2, FLT3, RET, others? | 31 |
| Bayer-18 | 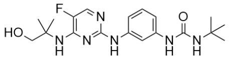 | Preclinical | TYK2, others? | 35 |
| ZM-39923 | 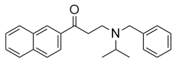 | Preclinical | **TGM2**, JAK1/3, EGFR | 36, 37 |
| Gandotinib | 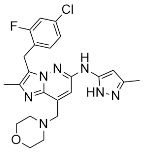 | Discontinued | JAK1/**2**, FLT3 | 38 |
| JAK3i | 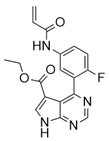 | Preclinical | **JAK3**, FLT3, TEC-family | 39 |
| NVP-BSK805 | 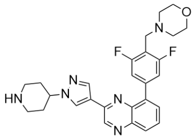 | Preclinical | **JAK2** | 40 |
| XL019 | 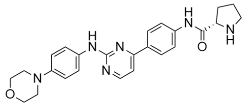 | Preclinical | **JAK2** | 41 |
| JAK2 inhibitor 13 | 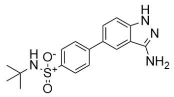 | Preclinical | JAK2, others? | 42 |
| TCS 21311 | 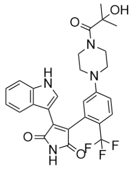 | Preclinical | **JAK3**, GSK-3β, PKCα and PKCθ | 43 |

| Compound | Structure | Status | Targets | References |
|---|---|---|---|---|
| Curvularin | 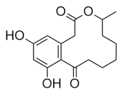 | Preclinical | JAK1/2, others? | 44 |
| JANEX-1 | 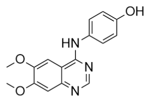 | Preclinical | **JAK3** | 45 |
| AG 490 | 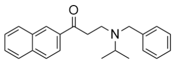 | Discontinued | **JAK2**/3, EGFR | 46, 47 |

### Supplementary References


(1) Parmentier, J. M.; Voss, J.; Graff, C.; Schwartz, A.; Argiriadi, M.; Friedman, M.; Camp, H. S.; Padley, R. J.; George, J. S.; Hyland, D.; Rosebraugh, M.; Wishart, N.; Olson, L.; Long, A. J. In Vitro and in Vivo Characterization of the JAK1 Selectivity of Upadacitinib (ABT-494). *BMC Rheumatol.* **2018**, *2*, 23. https://doi.org/10.1186/s41927-018-0031-x.

(2) Clark, J. D.; Flanagan, M. E.; Telliez, J. B. Discovery and Development of Janus Kinase (JAK) Inhibitors for Inflammatory Diseases. *Journal of Medicinal Chemistry*. American Chemical Society June 26, 2014, pp 5023–5038. https://doi.org/10.1021/jm401490p.

(3) Fridman, J. S.; Scherle, P. A.; Collins, R.; Burn, T. C.; Li, Y.; Li, J.; Covington, M. B.; Thomas, B.; Collier, P.; Favata, M. F.; Wen, X.; Shi, J.; McGee, R.; Haley, P. J.; Shepard, S.; Rodgers, J. D.; Yeleswaram, S.; Hollis, G.; Newton, R. C.; Metcalf, B.; Friedman, S. M.; Vaddi, K. Selective Inhibition of JAK1 and JAK2 Is Efficacious in Rodent Models of Arthritis: Preclinical Characterization of INCB028050. *J. Immunol.* **2010**, *184* (9), 5298–5307. https://doi.org/10.4049/jimmunol.0902819.

(4) Ito, M.; Yamazaki, S.; Yamagami, K.; Kuno, M.; Morita, Y.; Okuma, K.; Nakamura, K.; Chida, N.; Inami, M.; Inoue, T.; Shirakami, S.; Higashi, Y. A Novel JAK Inhibitor, Peficitinib, Demonstrates Potent Efficacy in a Rat Adjuvant-Induced Arthritis Model. *J. Pharmacol. Sci.* **2017**, *133* (1), 25–33. https://doi.org/10.1016/j.jphs.2016.12.001.

(5) Wernig, G.; Kharas, M. G.; Okabe, R.; Moore, S. A.; Leeman, D. S.; Cullen, D. E.; Gozo, M.; McDowell, E. P.; Levine, R. L.; Doukas, J.; Mak, C. C.; Noronha, G.; Martin, M.; Ko, Y. D.; Lee, B. H.; Soll, R. M.; Tefferi, A.; Hood, J. D.; Gilliland, D. G. Efficacy of TG101348, a Selective JAK2 Inhibitor, in Treatment of a Murine Model of JAK2V617F-Induced Polycythemia Vera. *Cancer Cell* **2008**, *13* (4), 311–320. https://doi.org/10.1016/j.ccr.2008.02.009.

(6) Elder, R. T.; Xu, X.; Williams, J. W.; Gong, H.; Finnegan, A.; Chong, A. S. F. The

Immunosuppressive Metabolite of Leflunomide, A77 1726, Affects Murine T Cells Through Two Biochemical Mechanisms. *J. Immunol.* **1997**, *159* (1), 22–27.

(7) Burke, J. R.; Cheng, L.; Gillooly, K. M.; Strnad, J.; Zupa-Fernandez, A.; Catlett, I. M.; Zhang, Y.; Heimrich, E. M.; McIntyre, K. W.; Cunningham, M. D.; Carman, J. A.; Zhou, X.; Banas, D.; Chaudhry, C.; Li, S.; D'Arienzo, C.; Chimalakonda, A.; Yang, X. X.; Xie, J. H.; Pang, J.; Zhao, Q.; Rose, S. M.; Huang, J.; Moslin, R. M.; Wrobleski, S. T.; Weinstein, D. S.; Salter-Cid, L. M. Autoimmune Pathways in Mice and Humans Are Blocked by Pharmacological Stabilization of the TYK2 Pseudokinase Domain. *Sci. Transl. Med.* **2019**, *11* (502). https://doi.org/10.1126/scitranslmed.aaw1736.

(8) Gu, Y. J.; Sun, W. Y.; Zhang, S.; Li, X. R.; Wei, W. Targeted Blockade of JAK/STAT3 Signaling Inhibits Proliferation, Migration and Collagen Production as Well as Inducing the Apoptosis of Hepatic Stellate Cells. *Int. J. Mol. Med.* **2016**, *38* (3), 903–911. https://doi.org/10.3892/ijmm.2016.2692.

(9) Kettle, J. G.; Åstrand, A.; Catley, M.; Grimster, N. P.; Nilsson, M.; Su, Q.; Woessner, R. Inhibitors of JAK-Family Kinases: An Update on the Patent Literature 2013-2015, Part 1. *Expert Opinion on Therapeutic Patents*. Taylor and Francis Ltd February 1, 2017, pp 127–143. https://doi.org/10.1080/13543776.2017.1252753.

(10) Coffey, G.; Betz, A.; DeGuzman, F.; Pak, Y.; Inagaki, M.; Baker, D. C.; Hollenbach, S. J.; Pandey, A.; Sinha, U. The Novel Kinase Inhibitor PRT062070 (Cerdulatinib) Demonstrates Efficacy in Models of Autoimmunity and B-Cell Cancer. *J. Pharmacol. Exp. Ther.* **2014**, *351* (3), 538–548. https://doi.org/10.1124/jpet.114.218164.

(11) Singer, J. W.; Al-Fayoumi, S.; Ma, H.; Komrokji, R. S.; Mesa, R.; Verstovsek, S. Comprehensive Kinase Profile of Pacritinib, a Nonmyelosuppressive Janus Kinase 2 Inhibitor. *J. Exp. Pharmacol.* **2016**, *8*, 11–19. https://doi.org/10.2147/JEP.S110702.

(12) Bissonnette, R.; Maari, C.; Forman, S.; Bhatia, N.; Lee, M.; Fowler, J.; Tyring, S.; Pariser, D.; Sofen, H.; Dhawan, S.; Zook, M.; Zammit, D. J.; Usansky, H.; Denis, L.; Rao, N.; Song, T.; Pavel, A. B.; Guttman-Yassky, E. The Oral Janus Kinase/Spleen Tyrosine Kinase Inhibitor ASN002 Demonstrates Efficacy and Improves Associated Systemic Inflammation in Patients with Moderate-to-Severe Atopic Dermatitis: Results from a Randomized Double-Blind Placebo-Controlled Study. *Br. J. Dermatol.* **2019**, *181* (4), 733–742. https://doi.org/10.1111/bjd.17932.

(13) Fensome, A.; Ambler, C. M.; Arnold, E.; Banker, M. E.; Brown, M. F.; Chrencik, J.; Clark, J. D.; Dowty, M. E.; Efremov, I. V.; Flick, A.; Gerstenberger, B. S.; Gopalsamy, A.; Hayward, M. M.; Hegen, M.; Hollingshead, B. D.; Jussif, J.; Knafels, J. D.; Limburg, D. C.; Lin, D.; Lin, T. H.; Pierce, B. S.; Saiah, E.; Sharma, R.; Symanowicz, P. T.; Telliez, J. B.; Trujillo, J. I.; Vajdos, F. F.; Vincent, F.; Wan, Z. K.; Xing, L.; Yang, X.; Yang, X.; Zhang, L. Dual Inhibition of TYK2 and JAK1 for the Treatment of Autoimmune Diseases: Discovery of ((S)-2,2-Difluorocyclopropyl)((1 R,5 S)-3-(2-((1-Methyl-1 H-Pyrazol-4-Yl)Amino)Pyrimidin-4-Yl)-3,8-Diazabicyclo[3.2.1]Octan-8-Yl)Methanone (PF-06700841). *J. Med. Chem.* **2018**, *61* (19), 8597–8612. https://doi.org/10.1021/acs.jmedchem.8b00917.

(14) Vazquez, M. L.; Kaila, N.; Strohbach, J. W.; Trzupek, J. D.; Brown, M. F.; Flanagan, M. E.; Mitton-Fry, M. J.; Johnson, T. A.; Tenbrink, R. E.; Arnold, E. P.; Basak, A.; Heasley, S. E.; Kwon, S.; Langille, J.; Parikh, M. D.; Griffin, S. H.; Casavant, J. M.; Duclos, B. A.; Fenwick, A. E.; Harris, T. M.; Han, S.; Caspers, N.; Dowty, M. E.; Yang, X.; Banker, M. E.; Hegen, M.; Symanowicz, P. T.; Li, L.; Wang, L.; Lin, T. H.; Jussif, J.; Clark, J. D.; Telliez, J. B.; Robinson, R. P.; Unwalla, R. Identification of N-{cis-3-[Methyl(7H-Pyrrolo[2,3-d]Pyrimidin-4-Yl)Amino]Cyclobutyl}propane-1-Sulfonamide (PF-04965842): A Selective JAK1 Clinical Candidate for the Treatment of Autoimmune Diseases. *J. Med. Chem.* **2018**, *61* (3), 1130–1152. https://doi.org/10.1021/acs.jmedchem.7b01598.

(15) Pardanani, A.; Lasho, T.; Smith, G.; Burns, C. J.; Fantino, E.; Tefferi, A. CYT387, a Selective JAK1/JAK2 Inhibitor: In Vitro Assessment of Kinase Selectivity and Preclinical Studies Using Cell Lines and Primary Cells from Polycythemia Vera Patients. *Leukemia* **2009**, *23* (8), 1441–1445. https://doi.org/10.1038/leu.2009.50.

(16) Telliez, J. B.; Dowty, M. E.; Wang, L.; Jussif, J.; Lin, T.; Li, L.; Moy, E.; Balbo, P.; Li, W.; Zhao, Y.; Crouse, K.; Dickinson, C.; Symanowicz, P.; Hegen, M.; Banker, M. E.; Vincent, F.; Unwalla, R.; Liang, S.; Gilbert, A. M.; Brown, M. F.; Hayward, M.; Montgomery, J.; Yang, X.; Bauman, J.; Trujillo, J. I.; Casimiro-Garcia, A.; Vajdos, F. F.; Leung, L.; Geoghegan, K. F.; Quazi, A.; Xuan, D.; Jones, L.; Hett, E.; Wright, K.; Clark, J. D.; Thorarensen, A. Discovery of a JAK3-Selective Inhibitor: Functional Differentiation of JAK3-Selective Inhibition over Pan-JAK or JAK1-Selective Inhibition. *ACS Chem. Biol.* **2016**, *11* (12), 3442–3451. https://doi.org/10.1021/acschembio.6b00677.

(17) Ludbrook, V. J.; Hicks, K. J.; Hanrott, K. E.; Patel, J. S.; Binks, M. H.; Wyres, M. R.; Watson, J.; Wilson, P.; Simeoni, M.; Schifano, L. A.; Reich, K.; Griffiths, C. E. M. Investigation of Selective JAK1 Inhibitor GSK2586184 for the Treatment of Psoriasis in a Randomized Placebo-Controlled Phase IIa Study. *Br. J. Dermatol.* **2016**, *174* (5), 985–995. https://doi.org/10.1111/bjd.14399.

(18) Wan, H.; Schroeder, G. M.; Hart, A. C.; Inghrim, J.; Grebinski, J.; Tokarski, J. S.; Lorenzi, M. V.; You, D.; Mcdevitt, T.; Penhallow, B.; Vuppugalla, R.; Zhang, Y.; Gu, X.; Iyer, R.; Lombardo, L. J.; Trainor, G. L.; Ruepp, S.; Lippy, J.; Blat, Y.; Sack, J. S.; Khan, J. A.; Stefanski, K.; Sleczka, B.; Mathur, A.; Sun, J. H.; Wong, M. K.; Wu, D. R.; Li, P.; Gupta, A.; Arunachalam, P. N.; Pragalathan, B.; Narayanan, S.; Nanjundaswamy, K. C.; Kuppusamy, P.; Purandare, A. V. Discovery of a Highly Selective JAK2 Inhibitor, BMS-911543, for the Treatment of Myeloproliferative Neoplasms. *ACS Med. Chem. Lett.* **2015**, *6* (8), 850–855. https://doi.org/10.1021/acsmedchemlett.5b00226.

(19) Wu, S. C.; Li, L. S.; Kopp, N.; Montero, J.; Chapuy, B.; Yoda, A.; Christie, A. L.; Liu, H.; Christodoulou, A.; vanBodegom, D.; vanderZwet, J.; Layer, J. V.; Tivey, T.; Lane, A. A.; Ryan, J. A.; Ng, S. Y.; DeAngelo, D. J.; Stone, R. M.; Steensma, D.; Wadleigh, M.; Harris, M.; Mandon, E.; Ebel, N.; Andraos, R.; Romanet, V.; Dölemeyer, A.; Sterker, D.; Zender, M.; Rodig, S. J.; Murakami, M.; Hofmann, F.; Kuo, F.; Eck, M. J.; Silverman, L. B.; Sallan, S. E.; Letai, A.; Baffert, F.; Vangrevelinghe, E.; Radimerski, T.; Gaul, C.; Weinstock, D. M. Activity of the Type II JAK2 Inhibitor CHZ868 in B Cell Acute Lymphoblastic Leukemia. *Cancer Cell* **2015**, *28* (1), 29–41. https://doi.org/10.1016/j.ccell.2015.06.005.

(20) Davis, M. I.; Hunt, J. P.; Herrgard, S.; Ciceri, P.; Wodicka, L. M.; Pallares, G.; Hocker,

M.; Treiber, D. K.; Zarrinkar, P. P. Comprehensive Analysis of Kinase Inhibitor Selectivity. *Nat. Biotechnol.* **2011**, *29* (11), 1046–1051. https://doi.org/10.1038/nbt.1990.

(21) Gozgit, J. M.; Bebernitz, G.; Patil, P.; Ye, M.; Parmentier, J.; Wu, J.; Su, N.; Wang, T.; Ioannidis, S.; Davies, A.; Huszar, D.; Zinda, M. Effects of the JAK2 Inhibitor, AZ960, on Pim/BAD/BCL-XL Survival Signaling in the Human JAK2 V617F Cell Line SET-2. *J. Biol. Chem.* **2008**, *283* (47), 32334–32343. https://doi.org/10.1074/jbc.M803813200.

(22) Thompson, J. E.; Cubbon, R. M.; Cummings, R. T.; Wicker, L. S.; Frankshun, R.; Cunningham, B. R.; Cameron, P. M.; Meinke, P. T.; Liverton, N.; Weng, Y.; DeMartino, J. A. Photochemical Preparation of a Pyridone Containing Tetracycle: A Jak Protein Kinase Inhibitor. *Bioorganic Med. Chem. Lett.* **2002**, *12* (8), 1219–1223. https://doi.org/10.1016/S0960-894X(02)00106-3.

(23) Anastassiadis, T.; Deacon, S. W.; Devarajan, K.; Ma, H.; Peterson, J. R. Comprehensive Assay of Kinase Catalytic Activity Reveals Features of Kinase Inhibitor Selectivity. *Nat. Biotechnol.* **2011**, *29* (11), 1039–1045. https://doi.org/10.1038/nbt.2017.

(24) Hedvat, M.; Huszar, D.; Herrmann, A.; Gozgit, J. M.; Schroeder, A.; Sheehy, A.; Buettner, R.; Proia, D.; Kowolik, C. M.; Xin, H.; Armstrong, B.; Bebernitz, G.; Weng, S.; Wang, L.; Ye, M.; McEachern, K.; Chen, H.; Morosini, D.; Bell, K.; Alimzhanov, M.; Ioannidis, S.; McCoon, P.; Cao, Z. A.; Yu, H.; Jove, R.; Zinda, M. The JAK2 Inhibitor AZD1480 Potently Blocks Stat3 Signaling and Oncogenesis in Solid Tumors. *Cancer Cell* **2009**, *16* (6), 487–497. https://doi.org/10.1016/j.ccr.2009.10.015.

(25) Goodacre, S. C.; Lai, Y.; Liang, J.; Magnuson, S. R.; Robarge, K. D.; Stanley, M. S.; Tsui, V. H.-W.; Williams, K.; Zhang, B.; Zhou, A. Janus Kinase Inhibitor Compounds and Methods. US-2010317643-A1, 2009.

(26) Pardanani, A.; Hood, J.; Lasho, T.; Levine, R. L.; Martin, M. B.; Noronha, G.; Finke, C.; Mak, C. C.; Mesa, R.; Zhu, H.; Soll, R.; Gilliland, D. G.; Tefferi, A. TG101209, a Small Molecule JAK2-Selective Kinase Inhibitor Potently Inhibits Myeloproliferative Disorder-Associated JAK2V617F and MPLW515L/K Mutations. *Leukemia* **2007**, *21* (8), 1658–1668. https://doi.org/10.1038/sj.leu.2404750.

(27) Ciceri, P.; Müller, S.; O'Mahony, A.; Fedorov, O.; Filippakopoulos, P.; Hunt, J. P.; Lasater, E. A.; Pallares, G.; Picaud, S.; Wells, C.; Martin, S.; Wodicka, L. M.; Shah, N. P.; Treiber, D. K.; Knapp, S. Dual Kinase-Bromodomain Inhibitors for Rationally Designed Polypharmacology. *Nat. Chem. Biol.* **2014**, *10* (4), 305–312. https://doi.org/10.1038/nchembio.1471.

(28) Stump, K. L.; Lu, L. D.; Dobrzanski, P.; Serdikoff, C.; Gingrich, D. E.; Dugan, B. J.; Angeles, T. S.; Albom, M. S.; Ator, M. A.; Dorsey, B. D.; Ruggeri, B. A.; Seavey, M. M. A Highly Selective, Orally Active Inhibitor of Janus Kinase 2, CEP-33779, Ablates Disease in Two Mouse Models of Rheumatoid Arthritis. *Arthritis Res. Ther.* **2011**, *13* (2). https://doi.org/10.1186/ar3329.

(29) Gonzales, A. J.; Bowman, J. W.; Fici, G. J.; Zhang, M.; Mann, D. W.; Mitton-Fry, M. Oclacitinib (APOQUEL®) Is a Novel Janus Kinase Inhibitor with Activity against Cytokines Involved in Allergy. *J. Vet. Pharmacol. Ther.* **2014**, *37* (4), 317–324. https://doi.org/10.1111/jvp.12101.

(30) Works, M. G.; Yin, F.; Yin, C. C.; Yiu, Y.; Shew, K.; Tran, T.-T.; Dunlap, N.; Lam, J.; Mitchell, T.; Reader, J.; Stein, P. L.; D'Andrea, A. Inhibition of TYK2 and JAK1 Ameliorates Imiquimod-Induced Psoriasis-like Dermatitis by Inhibiting IL-22 and the IL-23/IL-17 Axis. *J. Immunol.* **2014**, *193* (7), 3278–3287. https://doi.org/10.4049/jimmunol.1400205.

(31) Noronha, G.; Mak, C. C.; Cao, J.; Renick, J.; McPherson, A.; Zeng, B.; Pathak, V. P.; Lohse, D. L.; Hood, J. D.; Soll, R. M. Bi-Aryl Meta-Pyrimidine Inhibitors of Kinases, 2005.

(32) Blaskovich, M. A.; Sun, J.; Cantor, A.; Turkson, J.; Jove, R.; Sebti, S. M. Discovery of JSI-124 (Cucurbitacin I), a Selective Janus Kinase/Signal Transducer and Activator of Transcription 3 Signaling Pathway Inhibitor with Potent Antitumor Activity against Human and Murine Cancer Cells in Mice. *Cancer Res.* **2003**.

(33) Malerich, J. P.; Lam, J. S.; Hart, B.; Fine, R. M.; Klebansky, B.; Tanga, M. J.; D'Andrea, A. Diamino-1,2,4-Triazole Derivatives Are Selective Inhibitors of TYK2 and JAK1 over JAK2 and JAK3. *Bioorganic Med. Chem. Lett.* **2010**, *20* (24), 7454–7457. https://doi.org/10.1016/j.bmcl.2010.10.026.

(34) Kapuria, V.; Peterson, L. F.; Fang, D.; Bornmann, W. G.; Talpaz, M.; Donato, N. J. Deubiquitinase Inhibition by Small-Molecule WP1130 Triggers Aggresome Formation and Tumor Cell Apoptosis. *Cancer Res.* **2010**, *70* (22), 9265–9276. https://doi.org/10.1158/0008-5472.CAN-10-1530.

(35) Bömer, U. D.; Bonin, A. D.; Bothe, U. D.; Buchmann, B. D.; Eis, K. D. New Phenyl-Pyrimidin-2-Yl-Amine Compounds Are Tyrosine Kinase 2 Inhibitors Useful for Treating e.g. Rheumatoid Arthritis, Crohn's Disease, Asthma, Multiple Sclerosis, Adult Respiratory Distress Syndrome, Allergic Alveolitis and Uveitis, 2009.

(36) Brown, G. R.; Bamford, A. M.; Bowyer, J.; James, D. S.; Rankine, N.; Tang, E.; Torr, V.; Culbert, E. J. Naphthyl Ketones: A New Class of Janus Kinase 3 Inhibitors. *Bioorganic Med. Chem. Lett.* **2000**, *10* (6), 575–579. https://doi.org/10.1016/S0960-894X(00)00051-2.

(37) Lai, T. S.; Liu, Y.; Tucker, T.; Daniel, K. R.; Sane, D. C.; Toone, E.; Burke, J. R.; Strittmatter, W. J.; Greenberg, C. S. Identification of Chemical Inhibitors to Human Tissue Transglutaminase by Screening Existing Drug Libraries. *Chem. Biol.* **2008**, *15* (9), 969–978. https://doi.org/10.1016/j.chembiol.2008.07.015.

(38) Ma, L.; Clayton, J. R.; Walgren, R. A.; Zhao, B.; Evans, R. J.; Smith, M. C.; Heinz-Taheny, K. M.; Kreklau, E. L.; Bloem, L.; Pitou, C.; Shen, W.; Strelow, J. M.; Halstead, C.; Rempala, M. E.; Parthasarathy, S.; Gillig, J. R.; Heinz, L. J.; Pei, H.; Wang, Y.; Stancato, L. F.; Dowless, M. S.; Iversen, P. W.; Burkholder, T. P. Discovery and Characterization of LY2784544, a Small-Molecule Tyrosine Kinase Inhibitor of JAK2V617F. *Blood Cancer J.* **2013**, *3* (4). https://doi.org/10.1038/bcj.2013.6.

(39) Tan, L.; Akahane, K.; McNally, R.; Reyskens, K. M. S. E.; Ficarro, S. B.; Liu, S.; Herter-

Sprie, G. S.; Koyama, S.; Pattison, M. J.; Labella, K.; Johannessen, L.; Akbay, E. A.; Wong, K. K.; Frank, D. A.; Marto, J. A.; Look, T. A.; Arthur, J. S. C.; Eck, M. J.; Gray, N. S. Development of Selective Covalent Janus Kinase 3 Inhibitors. *J. Med. Chem.* **2015**, *58* (16), 6589–6606. https://doi.org/10.1021/acs.jmedchem.5b00710.

(40) Baffert, F.; Régnier, C. H.; De Pover, A.; Pissot-Soldermann, C.; Tavares, G. A.; Blasco, F.; Brueggen, J.; Chène, P.; Drueckes, P.; Erdmann, D.; Furet, P.; Gerspacher, M.; Lang, M.; Ledieu, D.; Nolan, L.; Ruetz, S.; Trappe, J.; Vangrevelinghe, E.; Wartmann, M.; Wyder, L.; Hofmann, F.; Radimerski, T. Potent and Selective Inhibition of Polycythemia by the Quinoxaline JAK2 Inhibitor NVP-BSK805. *Mol. Cancer Ther.* **2010**, *9* (7), 1945–1955. https://doi.org/10.1158/1535-7163.MCT-10-0053.

(41) Forsyth, T.; Kearney, P. C.; Kim, B. G.; Johnson, H. W. B.; Aay, N.; Arcalas, A.; Brown, D. S.; Chan, V.; Chen, J.; Du, H.; Epshteyn, S.; Galan, A. A.; Huynh, T. P.; Ibrahim, M. A.; Kane, B.; Koltun, E. S.; Mann, G.; Meyr, L. E.; Lee, M. S.; Lewis, G. L.; Noguchi, R. T.; Pack, M.; Ridgway, B. H.; Shi, X.; Takeuchi, C. S.; Zu, P.; Leahy, J. W.; Nuss, J. M.; Aoyama, R.; Engst, S.; Gendreau, S. B.; Kassees, R.; Li, J.; Lin, S. H.; Martini, J. F.; Stout, T.; Tong, P.; Woolfrey, J.; Zhang, W.; Yu, P. SAR and in Vivo Evaluation of 4-Aryl-2-Aminoalkylpyrimidines as Potent and Selective Janus Kinase 2 (JAK2) Inhibitors. *Bioorganic Med. Chem. Lett.* **2012**, *22* (24), 7653–7658. https://doi.org/10.1016/j.bmcl.2012.10.007.

(42) Antonysamy, S.; Hirst, G.; Park, F.; Sprengeler, P.; Stappenbeck, F.; Steensma, R.; Wilson, M.; Wong, M. Fragment-Based Discovery of JAK-2 Inhibitors. *Bioorganic Med. Chem. Lett.* **2009**, *19* (1), 279–282. https://doi.org/10.1016/j.bmcl.2008.08.064.

(43) Thoma, G.; Nuninger, F.; Falchetto, R.; Hermes, E.; Tavares, G. A.; Vangrevelinghe, E.; Zerwes, H. G. Identification of a Potent Janus Kinase 3 Inhibitor with High Selectivity within the Janus Kinase Family. *J. Med. Chem.* **2011**, *54* (1), 284–288. https://doi.org/10.1021/jm101157q.

(44) Yao, Y.; Hausding, M.; Erkel, G.; Anke, T.; Förstermann, U.; Kleinert, H. Sporogen, S14-95, and S-Curvularin, Three Inhibitors of Human Inducible Nitric-Oxide Synthase Expression Isolated from Fungi. *Mol. Pharmacol.* **2003**, *63* (2), 383–391. https://doi.org/10.1124/mol.63.2.383.

(45) Sudbeck, E. A.; Liu, X. P.; Narla, R. K.; Mahajan, S.; Ghosh, S.; Mao, C.; Uckun, F. M. Structure-Based Design of Specific Inhibitors of Janus Kinase 3 as Apoptosis-Inducing Antileukemic Agents. *Clin. Cancer Res.* **1999**.

(46) Gazit, A.; Osherov, N.; Posner, I.; Yaish, P.; Poradosu, E.; Levitzki, A.; Gazit, A.; Gilon, C. Tyrphostins. 2. Heterocyclic and α-Substituted Benzylidenemalononitrile Tyrphostins as Potent Inhibitors of EGF Receptor and ErbB2/Neu Tyrosine Kinases. *J. Med. Chem.* **1991**, *34* (6), 1896–1907. https://doi.org/10.1021/jm00110a022.

(47) Wang, L. H.; Kirken, R. A.; Erwin, R. A.; Yu, C. R.; Farrar, W. L. JAK3, STAT, and MAPK Signaling Pathways as Novel Molecular Targets for the Tyrphostin AG-490 Regulation of IL-2-Mediated T Cell Response. *J. Immunol.* **1999**.

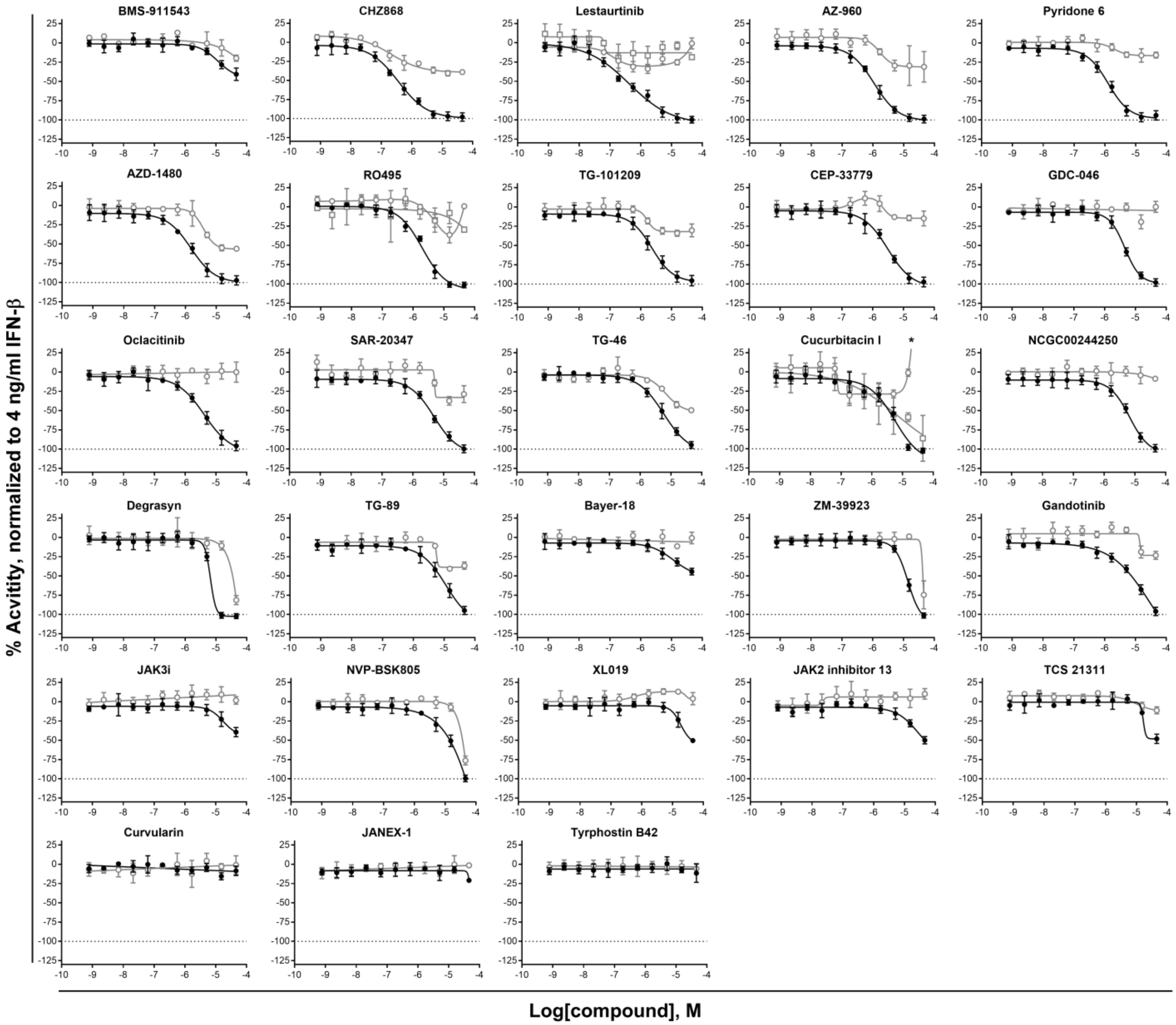


**Supplemental Figure 1.** CRCs for additional jakinibs. HLA-B HiBit clone AB-5 cells plated in 1536 well plates and treated for 24 hours with 4.0 ng/ml IFN-β plus 11-point, 1:3 titrations of compounds. Black circles – HiBit, open gray circles – CTG, open gray squares - CTF. For HiBit efficacy assays, data was normalized within each plate to 4.0 ng/ml IFN-β = 0% and DMSO alone = -100% activity. For CTG and CTF cytotoxicity assays, data was normalized within each plate to 4.0 ng/ml IFN-β = 0% and 92 uM Digitonin = -100% activity. *Highest concentration data point for curcubitacin I CTF was 250 ± 7%
